# BART Cancer: a web resource for transcriptional regulators in cancer genomes

**DOI:** 10.1101/2020.12.17.423327

**Authors:** Zachary V. Thomas, Zhenjia Wang, Chongzhi Zang

## Abstract

Dysregulation of gene expression plays an important role in cancer development. Identifying transcriptional regulators, including transcription factors and chromatin regulators, that drive the oncogenic gene expression program is a critical task in cancer research. Genomic profiles of active transcriptional regulators from primary cancer samples are limited in the public domain. Here we present BART Cancer (bartcancer.org), an interactive web resource database to display the putative transcriptional regulators that are responsible for differentially regulated genes in 15 different cancer types in The Cancer Genome Atlas (TCGA). BART Cancer integrates over 10,000 gene expression profiling RNA-seq datasets from TCGA with over 7,000 ChIP-seq datasets from the Cistrome Data Browser database and the Gene Expression Omnibus (GEO). BART Cancer uses Binding Analysis for Regulation of Transcription (BART) for predicting the transcriptional regulators from the differentially expressed genes in cancer samples compared to normal samples. BART Cancer also displays the activities of over 900 transcriptional regulators across cancer types, by integrating computational prediction results from BART and the Cistrome Cancer database. Focusing on transcriptional regulator activities in human cancers, BART Cancer can provide unique insights into epigenetics and transcriptional regulation in cancer, and is a useful data resource for genomics and cancer research communities.

## INTRODUCTION

Cancer and its progression arise from dysregulation of gene expression (1,2). The identification of transcriptional regulators (TRs), including transcription factors (TFs) and chromatin regulators (CRs), that control oncogenic gene expression program in each cancer type is an essential task for cancer research as these regulators could be targets for novel therapies. The Cancer Genome Atlas (TCGA) consortium has generated more than 10,000 RNA-seq datasets for tumor and normal samples for over 30 cancer types, as one of the largest resources for gene expression profiles in human cancers (3). Knowing what TRs regulate the genes with differential expression between tumor and normal samples can help better understand functional gene regulatory networks in each cancer type. However, TCGA did not produce ChIP-seq data due to limited cell numbers for most primary tumor samples. Therefore, computational prediction of functional TRs from differential gene expression data from TCGA will be useful in understanding cancer transcriptional regulation. Such efforts include the Cistrome Cancer web resource, which uses a comprehensive computational modeling approach to integrate TCGA data with publicly available chromatin profiling ChIP-seq data and reports the results on predicted TF targets and putative enhancer profiles in each cancer type (4). TF activities in Cistrome Cancer were inferred primarily based on the expression of the TF genes from TCGA and correlations with their putative target genes. The direct binding profiles of TFs from ChIP-seq data and their association with the putative enhancer profiles were not utilized in the model. Additionally, Cistrome Cancer focused on 575 factors, while growing ChIP-seq data from the public domain have covered more than 900 TRs (5–7), calling for an expansion of the existing database.

We previously developed Binding Analysis for Regulation of Transcription (BART), a computational method for inference of functional TRs based a target gene set as input (8). BART employs over 7,000 binding profiles for over 900 TRs to build the inference model for human, and has good performance in identifying the true TRs using differentially expressed genes generated from TR knock-down/knock-out experiments (9). BART can be utilized for inferring functional TRs from differentially expressed genes derive from TCGA data, providing a new approach for integration of tumor molecular profiling data and public ChIP-seq data. By using BART on both up and downregulated genes in cancer compared to normal, over 900 TRs can be reported based on their ranks of predicted activity in each cancer type. This allows us to identify not only regulators that promote tumorigenesis (from upregulated genes) but also regulators that may act as tumor suppressors (from downregulated genes). Here, we present BART Cancer, an interactive web resource database to report the putative transcriptional regulators that are responsible for differentially expressed genes in 15 different cancer types in TCGA, and multiple activity measures of over 900 transcriptional regulators across these 15 cancer types. An overview of the workflow can be seen in Figure 1.

**Figure 1.**
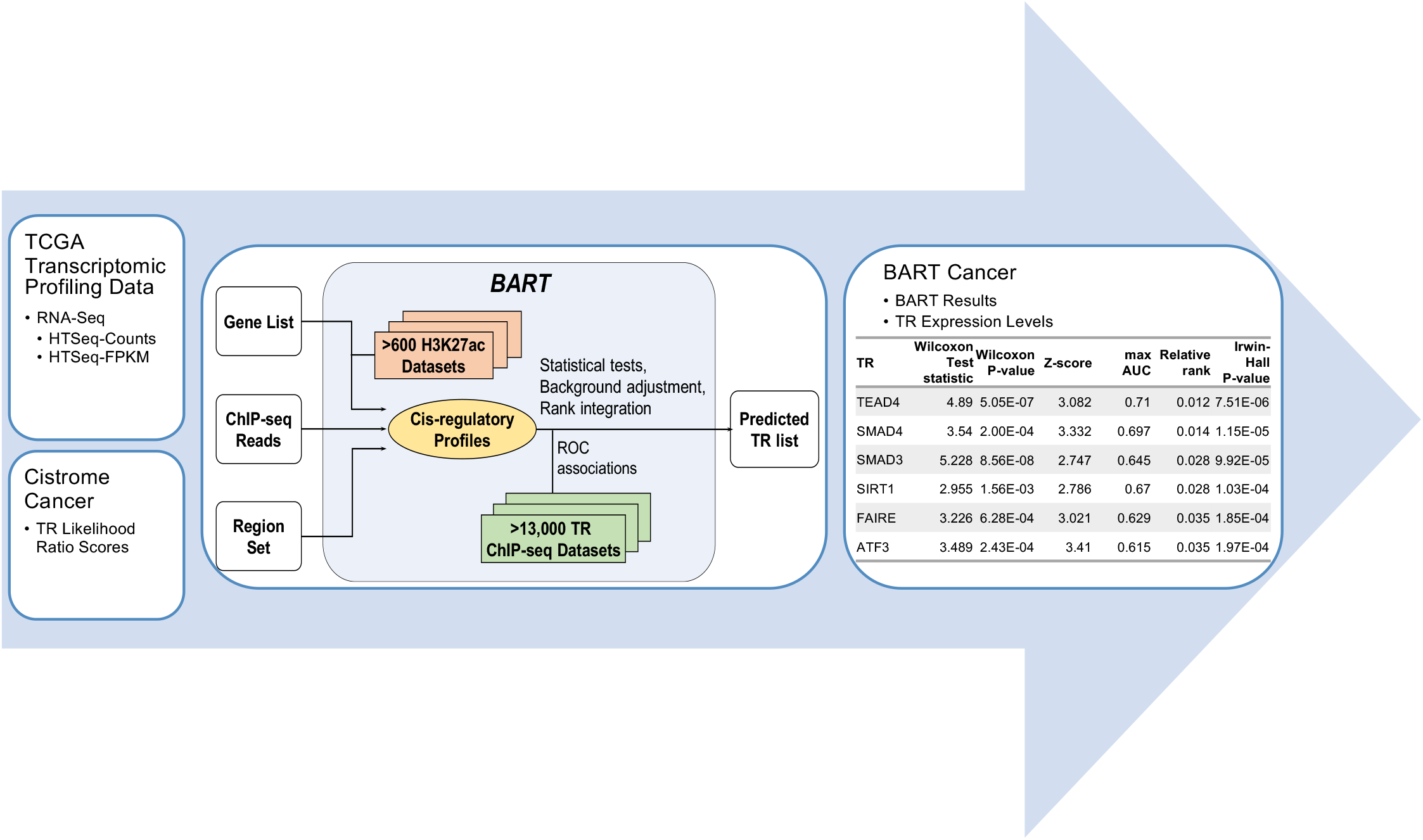
Schematic Representation of BART Cancer Workflow. RNA sequencing profiles were collected from TCGA and used to determine differentially expressed genes. These gene lists were then used as input to BART to get a putative transcriptional regulator (TR) list. This TR list, combined with Cistrome Cancer likelihood ratio scores, are displayed in BART Cancer.

## MATERIALS AND METHODS

### TCGA data collection and processing

RNA-seq data from 31 cancer types were downloaded from TCGA. In order to obtain a robust set of gene expression profiles, normalized read count in FPKM data from over 10,000 cancer samples were clustered using K-means clustering. 31 clusters were retained and the clusters from unambiguous cancer type annotation were kept with the TCGA annotation. Misclassified samples (<100) were removed from other cancer type clusters. Clusters with similar expression patterns from multiple cancer types were merged, e.g., esophagus-stomach cancer (STES) was stomach adenocarcinoma (STAD) and esophageal carcinoma (ESCA) combined; colorectal cancer was colon adenocarcinoma (COAD) and rectum adenocarcinoma (READ) combined. Additionally, breast cancer (BRCA) was clustered into two separate clusters: BRCA_1 for mainly luminal breast cancer (764 of 833 are luminal) and BRCA_2 for mainly basal breast cancer (176 of 234 are basal), similar to the results in Cistrome Cancer (4). A cancer type was only included in BART Cancer if TCGA included both cancer genomic profiles and normal genomic profiles, leaving BART Cancer with 15 different reannotated cancer types. For each cancer type, both up- and down-regulated genes were identified using DESeq2 (10). Differentially expressed genes were defined as those having adjusted *p*-value less than 1e-7 and log2(Fold Change) great than one for up-regulated genes and less than −1 for down-regulated genes. Differentially expressed genes provided by Cistrome Cancer (4) were used to cross-reference the newly identified cancer up-regulated genes. Because Cistrome Cancer does not provide down-regulated genes, newly identified down-regulated genes were not cross-referenced against any sources. These genes were then used as input to BART to get the putative transcriptional regulators responsible for regulating each gene set.

### BART analysis for TR prediction

Binding Analysis for Regulation of Transcription (BART) is a bioinformatics tool for predicting functional transcriptional regulators that bind at cis-regulatory regions to regulate gene expression in human or mouse (8). BART leverages a large collection of over 7,000 human TR binding profiles and 5,000 mouse TR binding profiles from the Cistrome DB. The main output of BART is a list of putative TRs with multiple quantification scores. The top ranked regulators are predicted putative regulators that are associated with the input dataset.

### Database design and implementation

BART Cancer is designed using HTML5, CSS, and JavaScript technologies. The Tabulator library (http://tabulator.info/) is used to display the BART results tables as well as list TRs to select plots to display. All TR plots were generated in R with ggplot2. The results, figures, and differentially expressed genes are all freely available for download.

The web-based user interface provides ways for researchers to view, visualize, and download the provided data. By selecting a cancer type and a type of differential expression, i.e., down- or up-regulated in cancer versus normal samples, the user is able to view the BART results of running BART on those genes for the specified cancer type. Directly below the display table include links to download the table as well as supplementary data files. Also provided is the link to the BARTweb (11) help page to better understand the meaning of the BART results (Figure 2).

**Figure 2.**
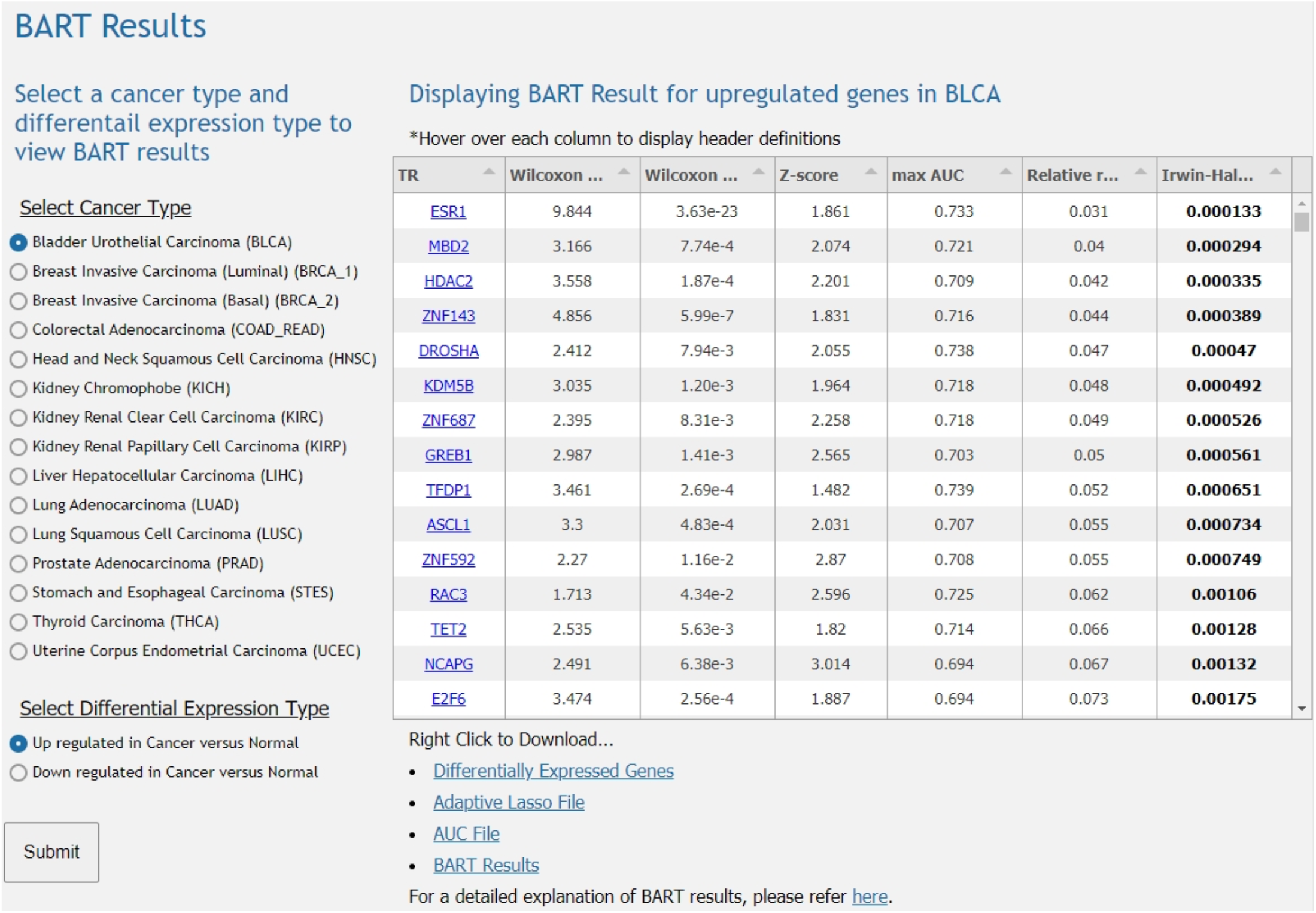
Example of BART Results Interface on BART Cancer. The radio button menus allow the user to select a reannotated TCGA cancer type and differential expression type. The BART results table then displays the list of putative transcriptional regulators for the cancer type and expression type selected. Below the table are links to download the tables and supplementary data files as well as a link to BARTweb for more information on BART results.

Below the BART results table module, TR specific plots are displayed. Each plot displays the expression level, BART rank, and the likelihood ratio score of a TR across all cancer types. TR gene’s average expression level was quantified as log(FPKM + 0.1) from TCGA RNA-seq data and ranked across the 15 cancer types. BART rank was quantified as 1-relative rank from the BART result. The likelihood ratio score was obtained from the Cistrome Cancer database, measuring the regulatory association of ChIP-seq data for this TR to the target genes predicted in Cistrome Cancer (4). This module uses a column of TR names which, when selected, change the figures displayed to the selected TR.

## RESULTS

BART Cancer web resource is created for users to browse putative TRs that are likely to regulate differential gene expression in a particular cancer and to visualize the relative activity of a TR across multiple cancer types. In order to validate that BART Cancer reports real functional TRs, we did a survey on the top three TRs predicted by BART from differentially expressed genes in each cancer type and cross-referenced with literature (Table 1).

**Table 1.** The top three transcriptional regulators from BART results on upregulated genes (left) and down regulated genes (right). The regulators in red are those that are discussed further as have literature-supported oncogenic activator or tumor suppressor functions. Cancer type abbreviations were following TCGA nomenclature.

| Upregulated Gene set BART Results |  |  |  | Downregulated Gene set BART Results |  |  |  |
| --- | --- | --- | --- | --- | --- | --- | --- |
| Cancer type | TR 1 | TR 2 | TR 3 | Cancer type | TR 1 | TR 2 | TR 3 |
| BLCA | ESR1 | MBD2 | HDAC2 | BLCA | MEF2B | NEUROG2 | MAML3 |
| BRCA1 | ESR1 | GREB1 | FOXA1 | BRCA1 | OTX2 | SOX2 | GATA4 |
| BRCA2 | MAX | POLR3D | TCF7L1 | BRCA2 | AR | GATA4 | REST |
| COAD_READ | TEAD4 | SMAD4 | SMAD3 | COAD_READ | REST | AR | MEF2B |
| HNSC | SMAD3 | SIRT1 | TEAD4 | HNSC | AR | FOXA1 | ESR1 |
| KICH | ERG | ESR1 | PR | KICH | HNF4A | RAD21 | HNF1A |
| KIRC | SITR1 | IKZF1 | TAL1 | KIRC | AR | HOXB13 | FOXA1 |
| KIRP | JUND | CEBPB | SMARCA4 | KIRP | MAML3 | RAD21 | GATA4 |
| LIHC | BRDU | ASCL1 | POLR3D | LIHC | GATA2 | TAL1 | CEBPB |
| LUAD | SIRT1 | ASCL1 | HNF4A | LUAD | GATA2 | TAL1 | GATA4 |
| LUSC | TP63 | GRHL2 | TP73 | LUSC | GATA2 | TAL1 | LYL1 |
| PRAD | AR | PIAS1 | FOXA1 | PRAD | OTX2 | SIX2 | TP63 |
| STES | TEAD4 | SMAD3 | YAP1 | STES | AR | REST | TBX5 |
| THCA | BRDU | TEAD4 | JUND | THCA | AR | OTX2 | NANOG |
| UCEC | GRHL2 | GREB1 | ASCL1 | UCEC | REST | CTCF | MAML3 |

### TRs for Up-regulated Genes

The TRs predicted from the up-regulated genes in cancer can be associated with oncogenic activation and promote tumorigenesis. Here we explore some of the top ranked TRs including histone deacetylase 2 (HDAC2), estrogen receptor 1 (ESR1), TEA domain family member 4 (TEAD4), achaete-scute homolog 1 (ASCL1), androgen receptor (AR), and Yes Associated Protein 1 (YAP1) in several cancer types.

In Bladder Urothelial Carcinoma (BLCA), HDAC2 ranked among the top three putative TRs. This class 1 deacetylase has been shown to be upregulated in bladder tumors (12). Additionally, a 2016 study showed that the inhibition of HDAC2 successfully inhibited cell proliferation and suggested that the inhibition of HDAC2 may be a promising therapy for urothelial carcinoma (13).

The top ranked putative regulator in luminal breast invasive carcinoma (BRCA luminal) was ESR1, which has been previously shown as a predictive biomarker of breast cancer. These results agree with previous studies on breast cancer that estrogen receptors and ESR1 mutations are markers of poor prognosis (14).

In colorectal adenocarcinoma (COAD & READ), TEAD4 has been shown to be a biomarker and tumorigenic promoter and has been identified as a possible target for therapies (15). The BART results reflect this as TEAD4 is the highest ranked regulator (Figure 3a).

**Figure 3.**
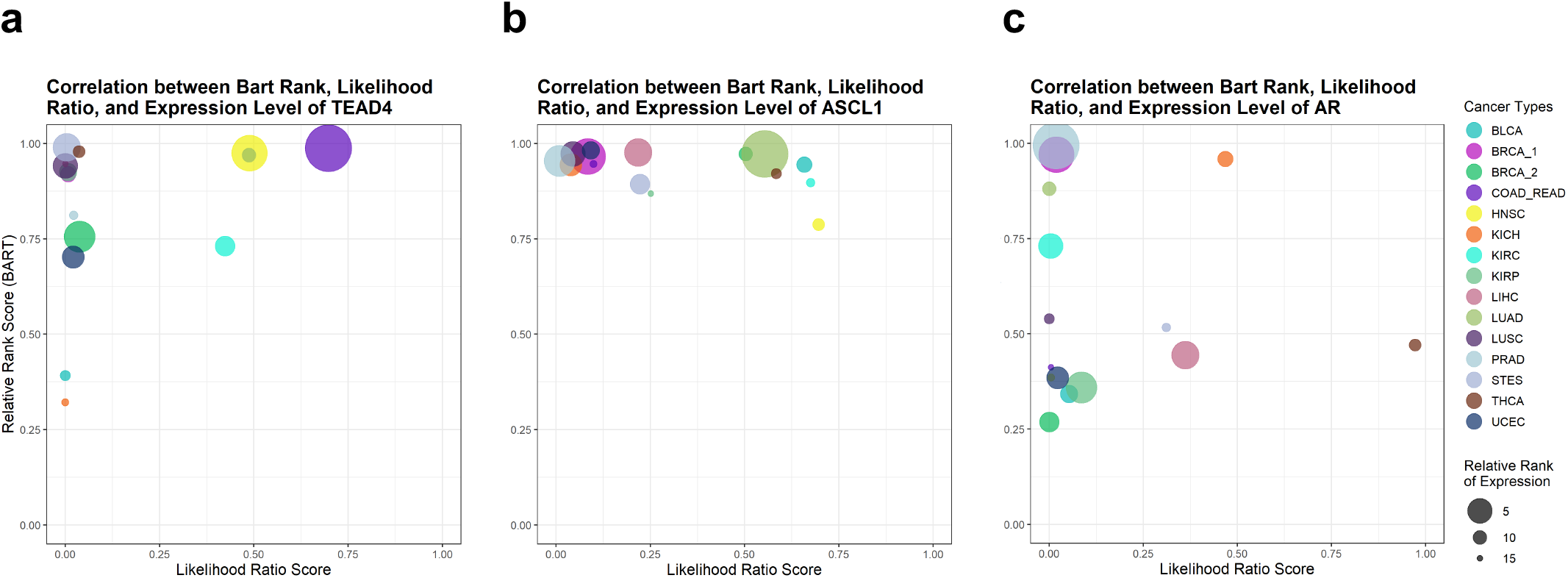
Integrative visualization of BART rank, Cistrome Cancer likelihood ratio score, and relative expression level of transcriptional regulators across cancer types. The bubble plots combine information from BART rank, Cistrome Cancer likelihood ratio score, and expression level. Each cancer type is plotted as a bubble on the 1 – relative rank vs likelihood ratio axes. The top right quadrant contains cancer types in which the transcriptional regulators ranked high in likelihood score and high in BART rank. The size of the bubble corresponds to the rank (1–15) of its relative expression level using log(FPKM + 0.1) from RNA-seq with the highest rank being the largest bubble, and the lowest rank being the smallest bubble. TEAD4 (**a**) and ASCL1 (**b**) show COAD_READ and LUAD in the top right quadrant, respectively, with the largest bubble indicating that they are likely oncogenic regulators in the corresponding cancer type. AR (**c**) shows PRAD as the largest bubble and highest BART rank. Although it does not rank high in Cistrome Cancer likelihood ratio score, it is still likely to be a functional transcription factor in PRAD.

In both lung cancers, lung adenocarcinoma (LUAD) and lung squamous cell carcinoma (LUSC), ASCL1 ranked among the top five putative regulators (Figure 3b). It has been demonstrated that ASCL1 is required for tumorigenesis in pulmonary cells (16). Additionally, some ASCL1 targets were ranked among the top twenty regulators including SOX2 whose overexpression has been clearly described in lung cancers (17).

In prostate adenocarcinoma (PRAD), the top ranked factor is AR (Figure 3c). This nuclear receptor has been shown to promote prostate cell growth and survival when bound to dihydrotestosterone (DHT). Thus, when there is an AR mutation or overexpression, prostate cell growth is directly impacted (18). For this reason, androgen deprivation therapy has been used to suppress prostate tumorigenesis (19).

YAP1 is a transcriptional co-activator and its activity with TEAD transcription factors have proliferative effects (20). BART results for up-regulated genes in stomach and esophageal carcinoma (STES) rank YAP1 and TEAD4 among the top three ranked putative TRs. This is supported by a 2016 study that has shown YAP1 knockdown suppresses esophageal carcinoma and proposes YAP1 to be a target for gene therapy (21).

### TRs for Down-regulated Genes

Transcriptional regulators predicted from down-regulated genes can be associated with tumor suppressor functions. Here we highlight six factors from the top three ranked TRs predicted from the down-regulated genes in several cancer types: GATA binding protein 4 (GATA4), Repressor element 1 (RE-1)-silencing transcription factor (REST), forkhead box A1 (FOXA1), ESR1, hepatocyte nuclear factor 4 alpha (HNF4A), and AR.

In breast cancer, one of the top three ranked transcriptional regulators for both basal and luminal breast cancer’s downregulated gene sets is GATA4. The dysregulation and aberrant expression of GATA4 has been correlated with tumor progression. While no direct tumor suppressive functions of GATA4 have been identified, it has been shown that restoration and even overexpression of GATA4 can impede the progression of breast tumors.

REST is ranked highly among many cancer’s BART results for downregulated genes. This result is not surprising as REST has previously been identified as a tumor suppressor from genetic screens (22). In breast cancer specifically, REST was identified to have significantly decreased expression in tumor compared to normal samples and REST knockdown in MCF-7 cells resulted in an increase in cell proliferation and reduced sensitivity to anticancer drugs (23). Similar results were found in colorectal cancer cell lines, further supporting its function as a tumor suppressor (24).

FOXA1 and ESR1, two of the top three ranked factors for head and neck cancer (HNSC) have been identified as tumor suppressors. First, second ranked FOXA1 was shown to suppress expression of miRNAs that were responsible for malignant behaviors in nasopharyngeal carcinoma, a subtype of head and neck squamous cell carcinoma (25). Second, third ranked ESR1 has been hypothesized to have tumor suppressive function in laryngeal squamous cell carcinoma (LSCC), another subcategory of head and neck cancer (26). It has also been shown that the removal of ESR1 results in a more aggressive LSCC and high ESR1 expression correlated to significantly improved survival (26).

HNF4A is identified as the top ranked regulator for kidney chromophobe (KICH). HNF4A has been associated with diabetes and has more recently been associated with reduction of proliferation in kidney cells, specifically renal cell carcinoma (27). HNF4A works by positively regulating genes that are responsible for regulating proliferation, specifically ABAT and ALDH6A1 (28, 29). By regulating these proliferation regulatory genes, HNF4A works as a tumor suppressor.

While AR was shown to be a oncogene, it has been shown to have the opposite effect in kidney chromophobe (KICH) and thyroid carcinoma (THCA). The loss of AR has been shown to correspond to increasing pathological grade of kidney caners (30). Further, it was shown that high expression of AR was associated with better survival rates supporting a tumor suppressor role for AR in kidney cancer (31). Likewise, decreased expression of AR in thyroid carcinoma was associated with poor prognosis and more aggressive tumor behavior (32).

## DISCUSSION

BART Cancer web resource provides information about computational model-predicted transcriptional regulators in 15 different cancer types, by integrating TCGA gene expression profiling data with publicly available ChIP-seq data for over 900 factors. It provides insights to cancer transcriptional regulation in two aspects. First, for researchers interested in a particular cancer type, BART Cancer reports putative transcription factors that either activate oncogenic gene expression or regulate tumor-suppressive gene expression. Depending on up- or downregulation of target genes and transcriptional activator or repressor function of the regulator, these putative regulators can have either oncogene or tumor-suppressor functions in this cancer. As many predicted transcription factors have been cross-referenced with literature, the novel factors in the prediction may guide further functional and mechanistic studies. Second, for researchers interested in a specific transcription factor or chromatin regulator, BART Cancer reports its relative activities across 15 cancer types and can give insights to which cancer types this factor might be functional in.

A few limitations of BART Cancer are worth noting. First, BART prediction was based on RNA-seq data from dozens to hundreds of TCGA samples and differentially expressed genes identified were shared across samples of each cancer type. The heterogeneity across samples were not taken into account. Also, cell heterogeneity could play a significant role especially in solid tumors, and the gene expression profiles could contain components from non-tumor cells such as tumor-infiltrating immune cells (4). As a result, BART predicted TRs might reflect cell composition changes rather than transcriptional regulation changes in cancer cells. Second, BART only utilizes publicly available ChIP-seq datasets for making predictions. There are over 1600 likely human transcription factors (33). Despite including more than 900 factors, there are still around 500 transcription factors that do not have high-quality ChIP-seq data and are absent in BART Cancer. The predicted TRs are far from complete. Third, discrepancies between BART rank and TR gene expression or likelihood ratio score were frequently found in TR plots. Possible reasons may include that differential transcription levels of a TR gene are not necessarily correlated with different TR activities. Likelihood ratio scores were based on putative TR target genes, whose accuracy could be limited because of the co-expression assumption and various ChIP-seq data qualities. Users should be aware of such caveats when interpreting BART Cancer data. In summary, despite of having certain limitations, BART Cancer can provide useful information in cancer transcriptional regulation and can be a valuable data resource for the cancer research community.

## AVAILABILITY

BART Cancer database resource is available at http://bartcancer.org/.

## ACKNOWLEDGEMENTS

The authors would like to thank Dr. Anindya Dutta for helpful discussions and Dr. Shenglin Mei for assistance on the Cistrome Cancer data.

## Author contributions

C.Z. conceived the study. Z.V.T. performed the analysis, interpreted the results, curated the database, and designed and implemented the web interface with supervision and assistance of Z.W. Z.V.T., Z.W. and C.Z. wrote the manuscript.

